# Differential effect of ethanol intoxication on peripheral markers of cerebral injury in murine blunt TBI

**DOI:** 10.1101/2020.09.18.303396

**Authors:** Zhenghui Li, Jin Zhang, Steffen Halbgebauer, Akila Chandrasekar, Rida Rehman, Albert Ludolph, Tobias Boeckers, Markus Huber-Lang, Markus Otto, Francesco Roselli, Florian olde Heuvel

## Abstract

Blood-based biomarkers have proven to be a reliable measure of traumatic brain injury (TBI) severity and outcome, in both murine models and patients. In particular, neuron-specific enolase (NSE) and neurofilament light (NFL) have been investigated in the clinical setting post injury. Ethanol intoxication (EI) remains a significant comorbidity in TBI, with 30-40% of patients having a positive blood alcohol level (BAC) post TBI. The effect of ethanol on blood-based biomarkers on the prognosis and diagnosis of TBI remain unclear. In this study, we investigated the effect of EI on NSE and NFL and their correlation with blood-brain barrier (BBB) integrity in a murine model of TBI. We have used ultra-sensitive single molecule array technology (SIMOA) and ELISA methods to measure NFL, NSE and Claudin-5 concentrations in plasma 3h post TBI. We showed that both NFL and NSE were increased 3h post TBI. However, ethanol blood concentrations only showed an inverse correlation with NSE, but not NFL. Claudin-5 levels were increased post injury, but no difference was detected in EI. The Claudin-5 increase post TBI was correlated with NFL, but not with NSE. Thus, the data indicate that ethanol has a confined effect on biomarker release in the bloodstream and neuronal biomarkers reflect a different pathophysiology upon TBI.

## Introduction

Blood based biomarkers serve as a promising candidate to define neuronal and glial damage upon Traumatic Brain Injury (TBI; 1,2). In particular, serum neuron-specific enolase (NSE) have been found to be increased in head trauma in patients (3,4) and NSE elevation was inversely correlated with the Glasgow Coma Scale score (5). Serum NSE levels were demonstrated to be predictive of TBI outcome (4). Alternative to NSE, Neurofilament light (NFL), which is a marker for axonal damage and is upregulated in various neurodegenerative disorders (6,7), has been recently shown to be upregulated in blood and CSF samples of TBI patients (8,9).

Peripheral biomarkers of neuronal damage have been infrequently applied to experimental models of TBI, in particular in murine models. NSE has been used as a proxy for the loss of neuronal integrity (10,11); plasma levels of NFL have been used to monitor the progression of neurodegeneration in murine models of Alzheimer’s disease (12) and it has been recently extended to applications in murine TBI (13).

Despite promising data from human and mouse cohort studies, several aspects of the application of peripheral biomarkers in the diagnosis and prognosis of TBI remain unclear. In particular, it has not been explored how important comorbidities interfere with the levels of peripheral biomarkers. In this context, ethanol intoxication is known to be one of the most common comorbidity in TBI, with 30-40% of patients showing a positive blood alcohol concentration (BAC) (14). The effect of BAC on patients’ survival and recovery remains controversial: positive BAC at the time of TBI has been associated with a better prognosis (15,16,17,18,19), but neutral or detrimental effects are also reported (20,21,22). In experimental TBI settings, ethanol proved to decrease TBI-induced neuroinflammatory effects and neuronal deficits (23,24,25,26), although not in all studies (27,28). In the present study, we have explored the effects of ethanol intoxication on peripheral biomarkers associated with TBI in a post-hoc analysis of murine samples from previous studies. Our goals were to determine if ethanol intoxication would affect the rise in brain damage markers upon TBI, focusing on the neuronal damage indicators NSE and NFL and on the vascular integrity marker Claudin-5. Here we show an increase in plasma levels of neuronal biomarkers NSE and NFL upon TBI. Blood ethanol concentrations display an inverse correlation with NSE but not with NFL. Notably, NFL levels directly correlated with Claudin-5, but not with NSE. These findings suggest different modes of biomarker release to the bloodstream after injury and the selective effect of ethanol on a subset of biomarkers used inTBI.

## Material & Methods

### Animals, Traumatic brain injury model and Ethanol treatment

This is a post-hoc, hypothesis-driven analysis of blood samples obtained in the context of previous studies (23,24,25,26); the investigation of these samples has never been reported before and it has been undertaken, in accordance to the 3R-principle, to reduce the number of mice employed in animal experimental and to maximize the scientific output from animal sacrifice.

These experiments have been approved by the Regierungspräsidium Tübingen with animal license no. 1222, with successive integrations, and by the Ulm University animal experimentation oversight committee.

B6-SJL were bred locally (Ulm University) under standard husbandry conditions (24°C, 60-80% humidity, 12/12 light/dark cycle, with ad libitum access to food and water). Experimental traumatic brain injury was performed as previously reported (23,24,25,26). Briefly, after administration of buprenorphine (0.1mg/kg by subcutaneously injection) and under sevoflurane anesthesia (2,5% in 97,5% O2), male mice aged 60-90 days were subject to closed weight-drop TBI. Animals were then manually positioned in the weight-drop apparatus and TBI was delivered by a 333g impactor free-falling from a 2cm distance, targeting the parietal bone (29). Immediately after the experimental TBI, animals were administered 100% O2 and were monitored for the apnea time. For sham surgery, mice were subjected to the same procedures and treatments (anesthesia, skin opening and closure, handling, positioning in the TBI apparatus), but no trauma was delivered. Ethanol was administered by oral gavage: 400µl of 32% ethanol solution in saline were administered 30 min before the procedures. Four experimental groups were considered: saline-administered, subjected to sham surgery (saline-sham, SS), saline-administered, subjected to TBI (saline-TBI, ST), ethanol administered, subjected to sham surgery (ethanol-sham, ES), ethanol administered, subjected to TBI (ethanol-TBI; ST).

### Blood sampling and plasma preparation

Three hours after trauma, animals were subject to xylazine/ketamine terminal anesthesia; blood was collected by right ventricular puncture using a 1-ml syringe equipped with a 24G needle and quickly transferred to a vial containing EDTA as anticoagulant. From each mouse, 400-500µl of blood (200-300µl of serum) was collected. Plasma samples were prepared by centrifuging the EDTA vials for 5min at 800g at 4°C, the supernatant was collected and centrifuged again for 2min at 13.000g at 4°C. Plasma was aliquoted and stored at -80°C until use.

### Blood Ethanol assay

The ethanol blood assay was performed according to manufacturer’s instructions (Abcam). Briefly, plasma was diluted in ddH_2_O (500x for ethanol samples and 10x for the saline samples). Samples were added to a 96-well plate, a mastermix of ethanol probe, ethanol enzyme mix and ethanol assay buffer was added and incubated for 30min at 37°C. The optical density (OD) 450nm was measured by using a colorimetric detection (Fluostar Optima, BMG Labtech) and concentrations were calculated with the standard curve.

### SIMOA

Single molecule array quantification of plasma Neurofilament light chain (Quanterix) was performed as previously reported (30) and in agreement with manufacturer’s instructions. Mouse serum (5µl) was diluted 1:20 before the assay.

### ELISA

Enzyme-linked immuno-sorbent assays for Neuron-specific Enolase (NSE) and Claudin-5 were performed according to manufacturer’s instructions (mouse ENO2/NSE (CLIA) ELISA Kit, LSbio; mouse Claudin-5(ELISA Kit, Cusabio). Briefly, 100µl of diluted samples (NSE: 1:10; Claudin-5: 1:2; diluted in sample diluent) or standards were added to the well and incubated 90 min at 37°C, followed by aspiration of the liquid and directly followed by 100µl of 1x biotinylated detection antibody working solution for 1hour at 37°C. The wells were aspirated and then washed 3x by adding 350µl of washing buffer to each well for 2min at RT, after which 100µl of 1x HRP conjugate was added to the well and incubated 30min at 37°C. The wells were aspirated and washed 5x with 350µl of washing buffer for 2min each. Finally, 100µl of working substrate solution was added to each well and incubated for 5min at 37°C (for the Claudin-5 ELISA 50µl of stop solution was added) after which the relative light units (RLU) (for NSE) or OD at 450nm (for Claudin-5) were measured by using a microplate luminometer (Luminoscan Ascent, Thermo Scientific). Concentrations were calculated according to the standard curve.

### Data analysis

Statistical analysis was performed using Graphpad prism version 8 software. Grouped analysis for the ELISA and SIMOA data was performed by using the Kruskal-Wallis test with Dunn’s multiple correction. Correlation assays were performed between treatment groups and different analytes to assess the relationship. Analysis of covariance (ANCOVA) was undertaken by comparing linear regression slopes of treatment groups with Tukey multiple correction. Data was depicted in graphs as median, 25th to 75th percentile (box), min to max (whiskers) or correlation with linear regression. Data in the text were depicted as median (min to max). Statistical significance was set at p < 0.05.

## Results

### Blood Ethanol concentration is not affected by TBI after oral binge

First, we established the concentration of ethanol in plasma upon single “oral binge” with or without concomitant TBI. For this, mice were pretreated with saline or ethanol (5g/kg) 30 min before either TBI or sham surgery; ethanol concentrations were measured 3h after the trauma. Four experimental groups were therefore considered: saline-sham (SS; N=8), ethanol-sham-S (ES; N=14), saline TBII (ST; N=24) and ethanol-TBI (ET; N=17). The Kruskal-Wallis test showed a significant effect in ethanol concentrations between treatment groups (p < 0.0001; Fig 1). The post-hoc comparison (Dunn’s corrected) revealed a significant difference, predictably, between SS and ES (median-range: 8.9 (0 to 84.0) µmol vs 4208.9 (1279.1 to 7661.6) µmol; p = 0.0003; Fig 1) and between ST and ET (median-range 13.5 (0 to 291.0) µmol vs 3832.1 (2186.5 to 7753.9) µmol; p < 0.0001; Fig 1) but ES and ET showed a comparable plasma ethanol concentrations (p > 0.9999; Fig 1). These findings suggest that TBI does not per se affect the clearance of ethanol after “oral binge”.

**Figure 1:**
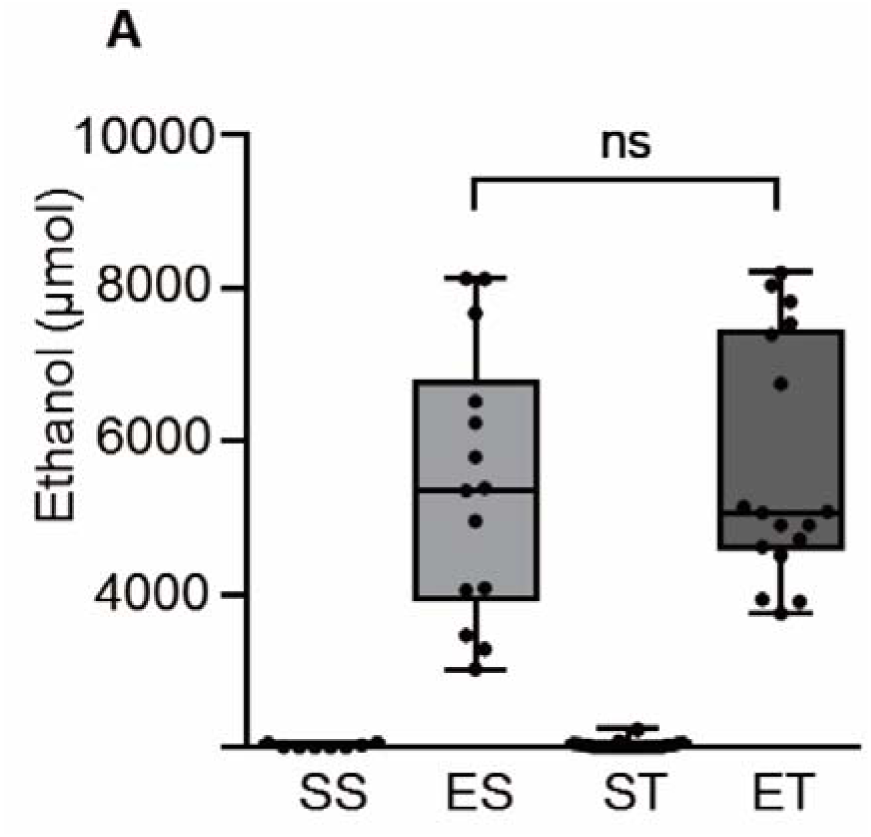
TBI does not affect ethanol concentration after binge administration. Blood alcohol concentration in blood plasma 3h after TBI, 4 treatment groups were used; saline-sham (SS), Ethanol-sham (ES), saline-TBI (ST) and ethanol-TBI (ET). Ethanol treatment showed a significant difference between treatment groups (p < 0.0001). Post-hoc analysis revealed a significant difference between SS and ES (p < 0.0003) and between ST and ET (p < 0.0001). However, ES and ET showed no difference (p > 0.9999). Boxplots represent median value, 25th to 75th percentile (box), min to max (whiskers), including individual data points. Group size: SS, N = 8; ES, N = 14; ST, N = 24; ET, N = 17. * : p < 0.05; ** : p < 0.01; *** : p < 0.001; **** : p < 0.0001.

### Ethanol intoxication blunts NSE but not NFL upregulation after TBI

Next we set out to investigate the effect of ethanol intoxication on two peripheral biomarkers of neuronal injury after TBI, namely NFL and NSE concentrations. For the NFL assessment, Kruskal-Wallis Test revealed a significant effect between treatment groups (p < 0.0001; Fig 2A). The post-hoc comparison (Dunn’s corrected) showed that ethanol alone, in absence of trauma, did not affect the NFL concentrations in plasma (median-range SS: 78.3 (35.8 to 226.2) pg/ml vs ES: 190.3 (20.9 to 1112.3) pg/ml; p > 0.9999; Fig 2A). TBI showed a significant upregulation in comparison to baseline (median-range SS: 78.3 (35.8 to 226.2) pg/ml vs ST: 969.4 (76.7 to 3276) pg/ml p = 0.0005; Fig 2A). Ethanol treatment prior to TBI showed only a trend towards lower concentrations in comparison to the ST group (ST: 969.4 (76.7 to 3276) pg/ml vs ET: 608.8 (238.5 to 1541.2) pg/ml; p > 0.9999; Fig 2A). Interestingly, concentrations of plasma ethanol were completely uncorrelated to NFL concentrations (in ET group R^2^ = 0.0170; p = 0.6306; Fig 2B).

**Figure 2:**
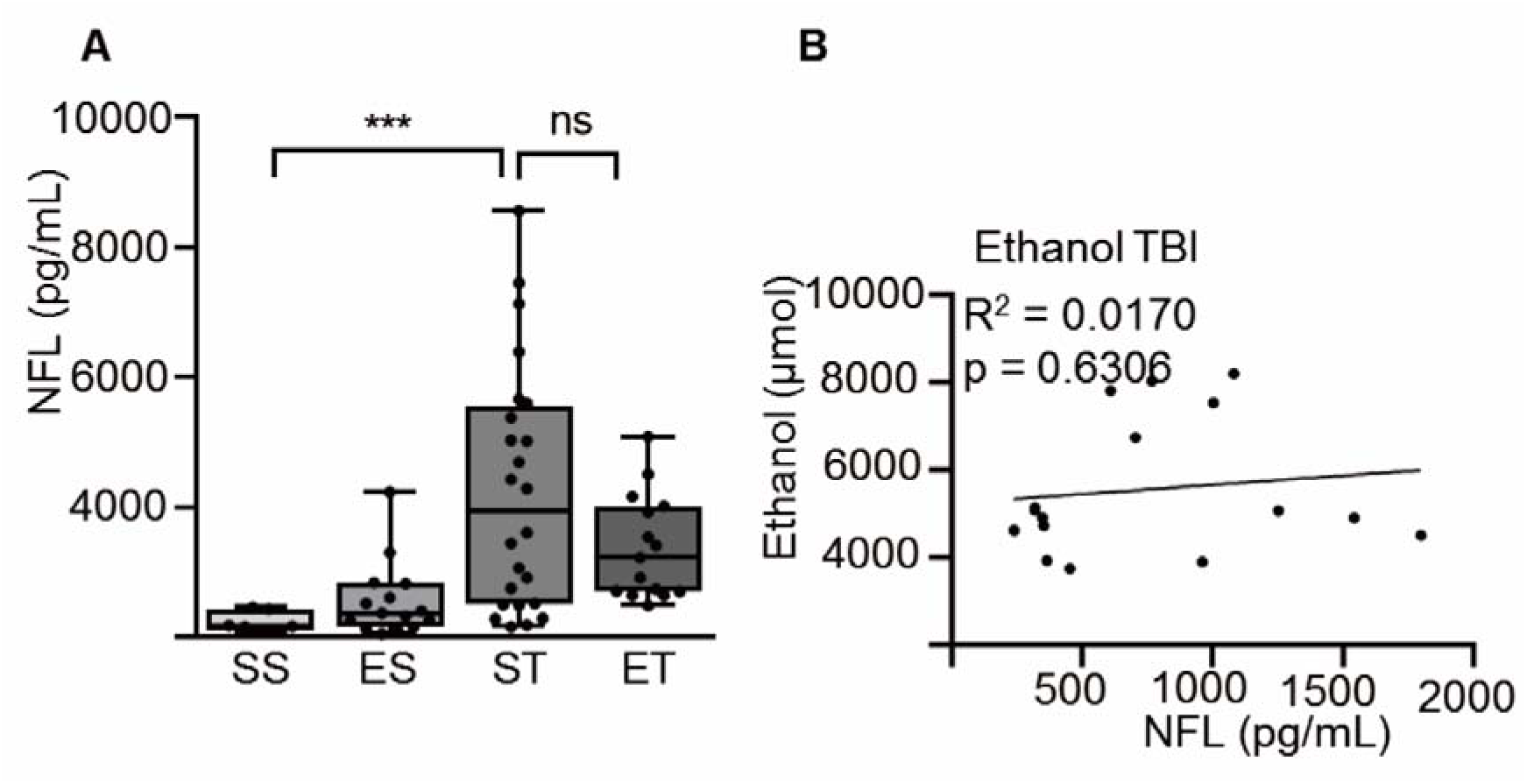
Ethanol does not affect NFL levels post TBI. NFL concentration was assessed 3h post TBI and Ethanol TBI, 4 treatment groups were used; saline-sham (SS), Ethanol-sham (ES), saline-TBI (ST) and ethanol-TBI (ET). A) Treatment groups showed significant differences in NFL concentration (p < 0.0001). Post-hoc analysis showed a significant difference between SS and ST (p = 0.0005) but no difference between ST and ET (p > 0.9999). B) Correlation analysis showed no significant relationship between NFL and ethanol in the ET group (R^2^ = 0.0170; p = 0.6306). Boxplots represent median value, 25th to 75th percentile (box), min to max (whiskers), including individual data points. Correlation with linear regression is shown. Group size: SS, N = 8; ES, N = 14; ST, N = 24; ET, N = 17. * : p < 0.05; ** : p < 0.01; *** : p < 0.001; **** : p < 0.0001.

Likewise, the Kruskal-Wallis Test applied to the NSE dataset revealed a significant effect between treatment groups (p = 0.0232; Fig 3A). The post-hoc comparison (Dunn’s corrected) revealed that ethanol alone showed no effect on NSE plasma concentrations (SS: 144.0 (98.6 to 291.5) vs ES: 191.9 (106.0 to 323.9); p > 0.9999; Fig 3A). TBI showed a significant effect in comparison to sham (median-range SS: 144.0 (98.6 to 291.5) vs ST: 215.7 (140.8 to 644.1); p = 0.0295; Fig 3A). However, ethanol treatment before TBI showed no significant difference between TBI on its own (median-range ST: 215.7 (140.8 to 644.1) pg/ml vs ET: 202.0 (65.9 to 380.5) pg/ml; p > 0.9999; Fig 3A), but a significant inverse correlation was found between ethanol concentrations and NSE concentrations in the ET group (R^2^ = 0.4069; p = 0.0078; Fig 3B)

**Figure 3:**
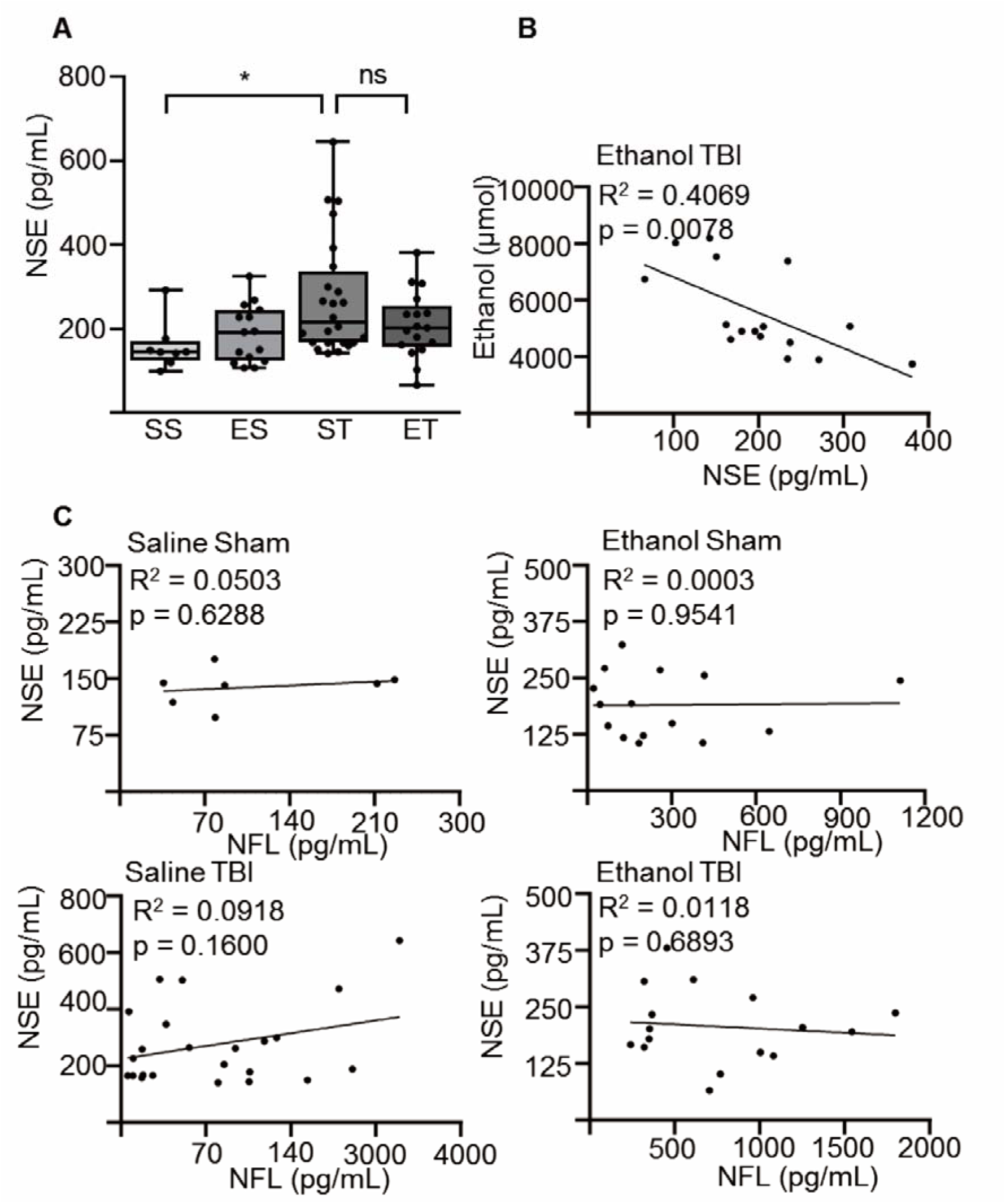
Ethanol decreases NSE levels post TBI. NSE concentration was assessed 3h post TBI and Ethanol TBI, 4 treatment groups were used; saline-sham (SS), Ethanol-sham (ES), saline-TBI (ST) and ethanol-TBI (ET). A) NSE concentration showed significant differences between treatment groups (p = 0.0232). Post-hoc analysis revealed a significant difference between SS and ST (p = 0.0295), but not between ST and ET (p > 0.9999). B) Correlation analysis revealed a significant inverse correlation between NSE and ethanol in the ET group (R^2^ = 0.4069; p < 0.0078). C) Slope comparison revealed no significant difference in the treatment groups between NFL and NSE (F = 0.4236; p = 0.7368), correlation within the groups showed no difference (NSE vs NFL SS: R^2^ = 0.0503; p = 0.6306, ES: R^2^ = 0.0003; p = 0.0954, ST: R^2^ = 0.0918; p = 0.1600, ET: R^2^ = 0.0118; p = 0.6893). Boxplots represent median value, 25th to 75th percentile (box), min to max (whiskers), including individual data points. Correlation with linear regression is shown. Group size: SS, N = 8; ES, N = 14; ST, N = 24; ET, N = 17. * : p < 0.05; ** : p < 0.01; *** : p < 0.001; **** : p < 0.0001.

Notably, the values of blood ethanol, NSE and NFL concentrations displayed a substantial variability, despite the use of inbred mouse lines and standardized equipment. Surprisingly, when we tested if the concentrations of the two neuronal damage markers were correlated, the ANCOVA revealed no difference between the slopes of the treatment groups (F = 0.4236; p = 0.7368) and correlation within the groups was poor (NSE vs NFL SS: R^2^ = 0.0503; p = 0.6306, ES: R^2^ = 0.0003; p = 0.9541, ST: R^2^ = 0.0918; p = 0.1600, ET: R^2^ = 0.0118; p = 0.6893; Fig 3C).

Thus, the amount of ethanol affects NSE plasma concentrations but not NFL plasma concentrations upon TBI, possibly indicating either an effect on the pathophysiology of the release of these biomarkers or an effect on their transport to the blood.

### NFL levels in the plasma is correlated to blood brain barrier disruption

Furthermore, we sought to identify if ethanol modified the disruption of blood-brain-barrier caused by TBI, and if such a disruption influenced NSE or NFL plasma concentrations. To this end we assessed Claudin-5 concentrations in our samples by ELISA. Plasma Claudin-5 concentrations have been repeatedly used as markers of loss of vascular integrity upon TBI (31,32). The Kruskal-Wallis Test revealed a significant difference between treatment groups (p = 0.0047; Fig 4A). The post-hoc analysis (Dunn’s corrected) showed no significant effect of ethanol by itself (median-range SS: 111.8 (46.1 to 168.7) pg/ml vs ES: 64.8 (0 to 180.0) pg/ml; p > 0.9999; Fig 4A). TBI resulted in an increase of plasma Claudin-5 concentrations compared to ES (ES: 64.8 (0 to 180.0) pg/ml vs ST: 161.2 (77.9 to 237.0) pg/ml; p = 0.0157; Fig 4A). However, ethanol did not alter the plasma Claudin-5 concentrations upregulated by TBI (ST: 161.2 (77.9 to 237.0) pg/ml vs ET: 178.0 (26.4 to 444.7) pg/ml; p > 0.9999; Fig 4A). When correlating the concentrations of Claudin-5 ET treatment group to the amount of ethanol in the same group, we identified no significant correlation between the concentrations of ethanol and the concentrations of Claudin-5 in the plasma (R^2^ = 0.0215; p = 0.5747; Fig 4B), suggesting that the amount of ethanol in the blood does not alter the disruption of blood-brain-barrier.

**Figure 4:**
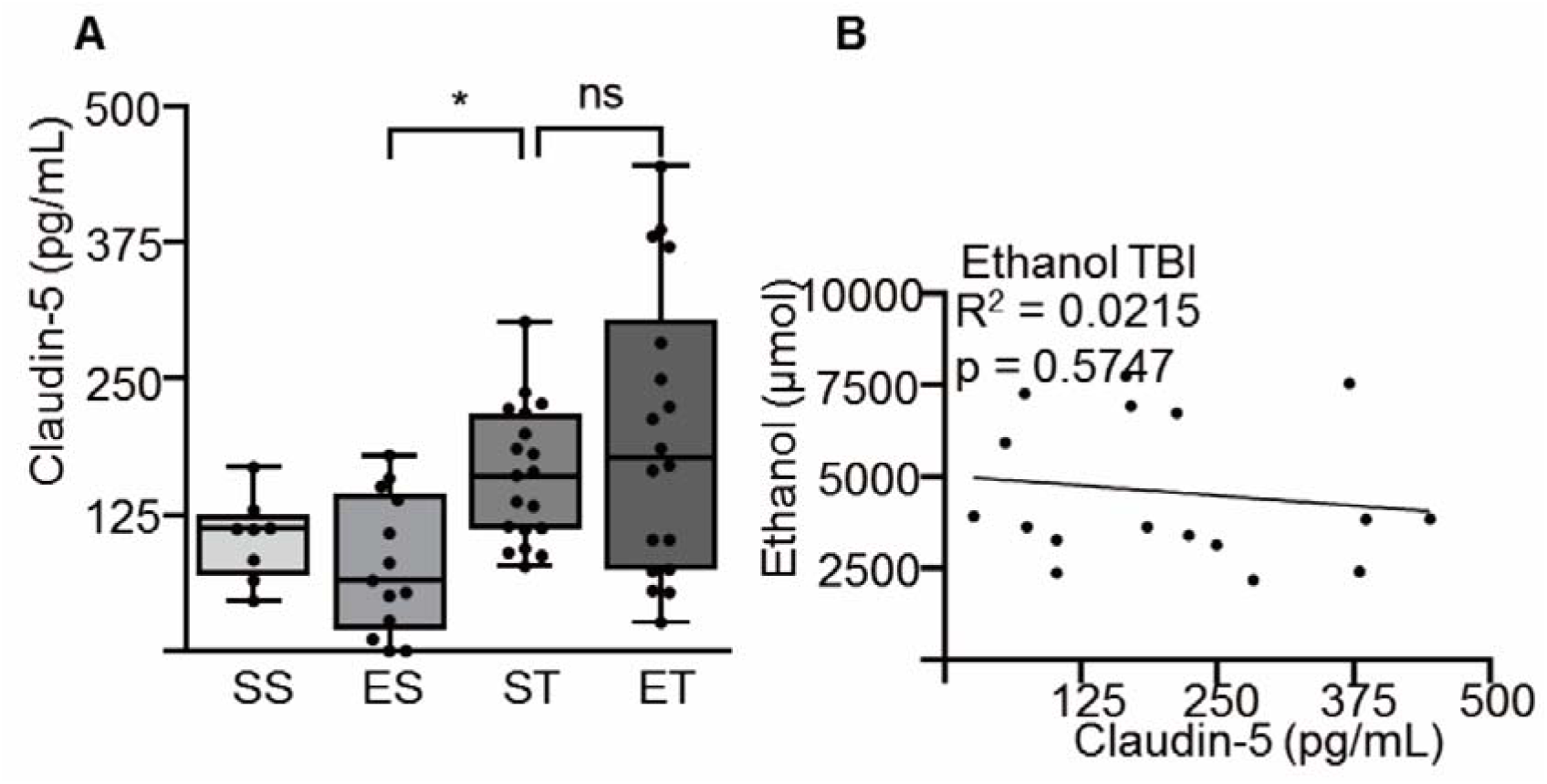
BBB disruption post TBI is unaffected by ethanol. Concentration of Claudin-5 was assessed in the blood 3h post TBI and ethanol TBI, 4 treatment groups were used; saline-sham (SS), Ethanol-sham (ES), saline-TBI (ST) and ethanol-TBI (ET). A) Treatment groups revealed significant differences in Claudin-5 concentration (p = 0.0047). Post-hoc analysis revealed significant differences between ES and ST (p = 0.0157) but no difference in ST and ET (p > 0.9999). B) Correlation analysis between Claudin-5 and ethanol showed no difference in the ET group (R^2^ = 0.0215; p = 0.5747). Boxplots represent median value, 25th to 75th percentile (box), min to max (whiskers), including individual data points. Correlation with linear regression is shown. Group size: SS, N = 8; ES, N = 14; ST, N = 24; ET, N = 17. * : p < 0.05; ** : p < 0.01; *** : p < 0.001; **** : p < 0.0001.

Next, we sought to identify the eventual correlation between the disruption of blood-brain-barrier and the concentrations of neuronal damage biomarkers. Analysis of covariance (ANCOVA) revealed no significant differences between the slopes of treatment groups (F = 0.9541; p = 0.4207). In fact, correlation of NSE and Claudin-5 concentrations was very poor in each of the four treatment groups (SS: R^2^ = 0.0379; p = 0.6757, ES: R^2^ = 0.0001; p = 0.9716, ST: R^2^ = 0.0470; p = 0.3455, ET: R^2^ = 0.1597; p = 0.1251; Fig 5A). However, when correlating NFL with Claudin-5 analysis of covariance (ANCOVA) showed a significant difference in treatment groups (F = 3.157; p = 0.0320); post-hoc analysis revealed a significant correlation in the ST group (R^2^ = 0.2545; p = 0.0276; Fig 5B), but not in the other treatment groups (SS: R^2^ = 0.5546; p = 0.0548, ES: R^2^ = 0.0512; p = 0.4571, ET: R^2^ = 0.0136; p = 0.6785; Fig 5B) and a significant difference between the ST and ET group (p = 0.0303). Thus, the severity of blood brain barrier disruption correlates with NFL concentrations but this is not true for NSE.

**Figure 5:**
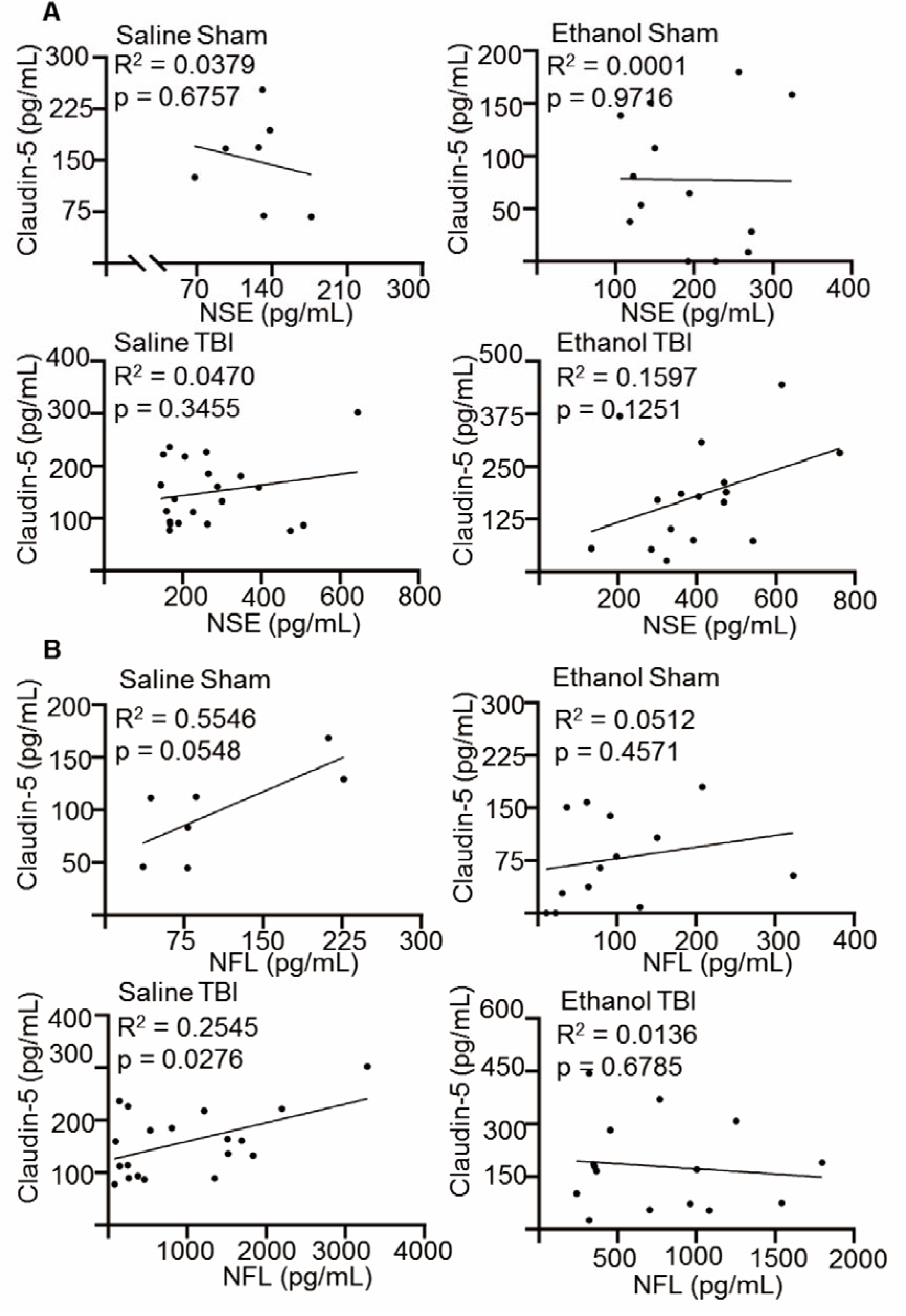
NFL levels are correlated to BBB disruption. Claudin-5 levels were correlated with both NSE and NFL concentration 3h post TBI and ethanol TBI, 4 treatment groups were used; saline-sham (SS), Ethanol-sham (ES), saline-TBI (ST) and ethanol-TBI (ET). A) Slope comparison revealed no significant difference in the treatment groups between NSE and Claudin-5 (F = 0.9541; p = 0.4207), correlation of NSE and Claudin-5 levels was very poor in each of the four treatment groups (SS: R^2^ = 0.0379; p = 0.6757, ES: R^2^ = 0.0001; p = 0.9716, ST: R^2^ = 0.0470; p = 0.3455, ET: R^2^ = 0.1597; p = 0.1251). B) Slope comparison revealed significant difference in the treatment groups between NFL and Claudin-5 (F = 3.157; p = 0.0320), correlation of NSE and Claudin-5 levels were significant in the ST group (R^2^ = 0.2545; p = 0.0276) but not in the other treatment groups (SS: R^2^ = 0.5546; p = 0.0548, ES: R^2^ = 0.0512; p = 0.4571, ET: R^2^ = 0.0136; p = 0.6785) and a significant difference between the ST and ET group (p = 0.0303). Correlation with linear regression is shown. Group size: SS, N = 8; ES, N = 14; ST, N = 24; ET, N = 17. * : p < 0.05; ** : p < 0.01; *** : p < 0.001; **** : p < 0.0001.

## Discussion

Here we have demonstrated that TBI induces the elevation of the peripheral damage related neuronal and vascular biomarkers (NSE, NFL and Claudin-5) and that ethanol appears to dose-dependently decrease NSE but not NFL or Claudin-5 concentrations in serum samples. Interestingly, even in absence of ethanol modulation, NFL serum concentrations correlated with the concentrations of Claudin-5, but not with NSE concentrations. Thus, our data suggest that i) ethanol intoxication reduces NSE concentrations, but not necessarily the overall acute burden of neurodegeneration as explored by NFL, and ii) concentrations of NFL, but not of NSE, correlates with the degree of disruption of the blood-brain-barrier (BBB).

Both NSE and NFL have been extensively investigated as markers of neuronal damage in TBI, displaying an overall significant elevation upon brain trauma (1,4,5,33,34). Nevertheless, the two markers do not appear to represent the same type of pathogenic process. Whereas NSE elevation peaks early after TBI, NFL levels keep increasing over several days (1) and their levels, although broadly elevated, do not always display a close correlation (1,35). This is not completely surprising when considering the different cell biology of the two markers: NFL is a constituent of the cytoskeleton and is highly enriched in large, myelinated axons, from where a small amount of NFL is released even in normal conditions (36,37). NSE is, on the other hand, an abundant soluble cytoplasmic protein expressed in neurons and in glial cells (38). Thus, whereas NFL is a better marker of axonal damage (39), NSE may reflect neuronal integrity. As a consequence, ethanol appears to be effective in limiting acute neuronal damage in TBI mouse models (in agreement with previous histological data; 10,24), but less effective (or ineffective) in protecting axonal tracts. These findings may be in accordance with activity-dependent mechanisms that may prevent neuronal degeneration, but may be unrelated to the dismantling of long-range axons. This observation may explain why ethanol intoxication in TBI patients appears to be protective especially in patients with penetrating injuries and much less in those with diffuse axonal injury (40).

Claudin-5 is a tight junction protein highly expressed in endothelial cells, where it contributes to the integrity of the BBB (41). Serum levels of Claudin-5 are considered a promising marker of BBB integrity (42,43). Although ethanol appears to decrease the inflammatory response induced by TBI (23), it does not affect the peripheral elevation in soluble Claudin-5 triggered by TBI. This finding is in agreement with the previous report of the lack of effect of ethanol on the downregulation of IL-25, a central regulator of BBB integrity, upon trauma (26). Thus, ethanol appears to have a restricted effect on peripheral biomarkers, reducing NSE without affecting NFL or Claudin-5 and thus possibly affecting neuronal survival, but not axonal damage and vascular integrity.

Interestingly, Claudin-5 concentrations directly correlated with NFL concentrations, but not with NSE concentrations upon trauma (with or without ethanol). This finding suggests that whereas the appearance of NFL in serum is facilitated by the increase in permeability of the BBB, serum NSE levels increase irrespective of BBB integrity. In fact, clearance of NSE from the CNS is only mildly related to BBB markers in TBI patients (44). It is tempting to speculate that other systems, such as the recently-identified glymphatic system, may be involved in the BBB-independent transfer of NSE from the brain to the blood. This finding further underscores how NFL and NSE may explore distinct aspects of the pathophysiology of TBI.

As a potential limitation of this study, it is worth noticing that we have used a relatively small number of biomarkers to investigate the effect of ethanol on biomarker prognosis. Various biomarkers might display distinct responses to ethanol upon TBI, like inflammatory mediators (45) or astrocyte-enriched proteins (46). The small number of biomarkers explored was due to the comparatively small volume of serum recovered from each mouse and the volume required by ELISA assays. A more widespread use of high-sensitivity SIMOA systems will allow in future the in-depth exploration of biomarkers of TBI in murine models. A second limitation may be constituted by the exploratory, retrospective design of this study; although operators and procedures were similar across several studies, unforeseen biases may contribute to the large variability of the NSE and NFL values. Therefore, this study should be considered as an entry point for further, dedicated, prospective studies using multiple biomarkers to address the effect of ethanol on specific cellular subpopulations and specific pathogenetic cascades.

## Acknowledgement

This work has been supported by the Deutsche Forschungsgemeinschaft as part of the Collaborative Research Center 1149 “Danger Response, Disturbance Factors and Regenerative Potential after Acute Trauma”. FR is also supported by the ERANET-NEURON initiative “External Insults to the Nervous System” as part of the MICRONET consortium and by the Baustein program of the Medical Faculty of Ulm University. Technical support by Thomas Lenk was highly appreciated.

